# Study on the efficacy of dsRNAs with increasing length in RNAi-based silencing of the *Fusarium CYP51* genes

**DOI:** 10.1101/824953

**Authors:** L Höfle, A Shrestha, B Werner, L Jelonek, A Koch

## Abstract

Previously, we have demonstrated that transgenic Arabidopsis and barley plants, expressing a 791 nucleotide (nt) dsRNA (CYP3RNA) that targets all three *CYP51* genes (*FgCYP51A, FgCYP51B, FgCYP51C*) in *Fusarium graminearum* (*Fg*), inhibited fungal infection via a process designated as host-induced gene silencing (HIGS). More recently, we have shown that spray applications of CYP3RNA also protect barley from fungal infection via a process termed spray-induced gene silencing (SIGS). Thus, RNAi technology may have the potential to revolutionize plant protection in agriculture. Therefore, successful field application will require optimization of RNAi design necessary to maximize the efficacy of the RNA silencing construct for making RNAi-based strategies a realistic and sustainable approach.

Previous studies indicate that silencing is correlated with the number of siRNAs generated from a dsRNA precursor. To prove the hypothesis that silencing efficiency is correlated with the number of siRNAs processed out of the dsRNA precursor, we tested in a HIGS and SIGS approach dsRNA precursors of increasing length ranging from 400 nt to 1500 nt to assess gene silencing efficiency of individual *FgCYP51* genes. Concerning HIGS-mediated disease control, we found that there is no significant correlation between the length of the dsRNA precursor and the reduction of *Fg* infection on CYP51-dsRNA expressing Arabidopsis plants. Importantly and in clear contrast to HIGS, we measured a decrease in SIGS-mediated *Fg* disease resistance that significantly correlates with the length of the dsRNA construct that was sprayed, indicating that the size of the dsRNA interferes with a sufficient uptake of dsRNAs by the fungus.

## Introduction

Crop plants are challenged by a multitude of different pathogens, insects, animals and weeds that cause many different plant diseases and constitute a constant threat to food supply. It is estimated that those organisms cause yield losses up to 40% of the global agricultural production what constitutes extremely high costs for our growing world population (Alexander et al., 2017; OERKE and Dehne, 2004). Besides the yield losses, mycotoxin contamination of foods and feedstuffs caused by phytopathogenic fungi, such as *Fusarium graminearum* (*Fg*), poses an almost intractable problem in agricultural production (Doll and Danicke, 2011). Current plant protection strategies rely on fungicide application for both *Fusarium* disease control and limitation of mycotoxin accumulation. The most commonly used fungicides are azoles, which target the cytochrome P450 sterol 14α-demethylase encoded by *FgCYP51* genes. Inhibition of CYP51 causes depletion of ergosterol which results in loss of membrane integrity followed by growth inhibition and death of fungal cells (Yoshida, 1988). However, as a consequence of continuous fungicide applications an increasing rate of azoles insensitivity was observed in several plant pathogenic fungi including *Fusarium* species (Becher et al., 2010; Spolti et al., 2014). Thus, the development of alternative control strategies has become one of the biggest challenges for plant pathologists in these days.

RNA interference (RNAi) is a conserved and integral aspect of gene regulation that utilizes small RNAs (sRNAs) to direct the silencing of gene expression at the transcriptional or posttranscriptional level. Over the last decade RNAi-based gene silencing has emerged as a powerful genetic tool for scientific research. It has been utilized in various fields of applied research, such as agriculture and human and veterinary medicine. The mechanism of RNAi-mediated plant protection strategies relies on the expression of the transgene that provokes the formation of a double stranded (ds)RNA precursor molecule *in planta*. This dsRNA subsequently triggers the plants RNAi machinery to process small interfering (si)RNAs that in case of an infection exhibit target gene silencing of a certain pathogen (Koch and Kogel 2014).

In a previous study we demonstrated that transgenic *Arabidopsis* and barley (*Hordeum vulgare*) plants, expressing a 791 nucleotide (nt) dsRNA (CYP3RNA) that targets all three *CYP51* genes (*FgCYP51A, FgCYP51B, FgCYP51C*) in *Fg*, inhibited fungal infection via a process designated as host-induced gene silencing (HIGS) (Koch et al., 2013; Nowara et al., 2010). Furthermore, HIGS has been shown to protect several different plants species against infection by nematodes (Shivakumara et al., 2017), insects (Abdellatef et al., 2015), bacteria (Walawage et al., 2013) and fungi (Koch et al., 2013) as well as invasion by parasitic plants (Alakonya et al., 2012).

While we already provided proof-of-concept that RNAi-based plant protection is an effective strategy for controlling diseases caused by devastating necrotrophic pathogens, the broad applicability of HIGS remains questionable due to fact that generation of genetically modified (GM) crops is time-consuming and weakly accepted in many European countries. Therefore, we established an RNAi-based non-GMO crop protection approach using direct spray applications of dsRNA to target pathogens. Recently, we have shown that spray applications of CYP3RNA also protect barley from fungal infection via a process termed spray-induced gene silencing (SIGS) (Koch et al., 2016). Our finding that inhibitory dsRNA is effective upon spray application is of a ground-breaking nature, and it represents significant progress to make RNAi-based approaches for plant protection scientifically and economically achievable. Whereas a great number of studies have been published on HIGS-mediated silencing of target genes in pathogenic microbes, silencing of such targets through exogenously applied dsRNA has been described only in a few studies (Koch et al., 2016; Mitter et al., 2017; Wang et al., 2016).

However, given the ease of dsRNA design, its high specificity, and applicability to diverse pathogens, the use of target-specific dsRNA as an anti-fungal agent offers an unprecedented potential as a new plant protection strategy. Therefore, successful field application will require optimization of RNAi design necessary to maximize the efficacy of the RNA silencing construct. Recently, we compared the efficiencies of HIGS and SIGS dsRNA delivery strategies to assess the activity of novel dsRNA species that were designed to target one or two *FgCYP51* genes (Höfle et al., 2018 in revision). Using barley as a cereal model, we found that dsRNA constructs targeting two *FgCYP51* genes inhibited fungal growth more efficient than single constructs, although both types of dsRNAs decreased fungal infections (Höfle et al., 2018 in revision). Based on these findings, we anticipate that constructs which target two genes in parallel were more efficient because the number of siRNAs derived from those double constructs are higher.

Previous studies led to the hypothesis that silencing is correlated with the number of siRNAs generated from a dsRNA precursor

Therefore, we tested in a HIGS and SIGS approach dsRNA precursors of increasing length ranging from 400 nt to 1500 nt to assess gene silencing efficiency of individual *FgCYP51* genes.

We found that SIGS efficiencies dependent on the length of the dsRNA that was sprayed, indicating that the size of the sprayed dsRNA interferes with a sufficient uptake mechanisms of the fungus. Interestingly, we found that HIGS-mediated disease resistance was independent of the length of the dsRNA constructs. Our findings suggest that HIGS and SIGS approaches differ concerning their mechanistic basis, thus leading to different silencing efficiencies and disease resistance phenotypes depending on RNAi construct design.

## Results and Discussion

### Host-induced gene silencing by CYP51-dsRNAs of different length confers resistance to *Fg* in transgenic *Arabidopsis*

To assess whether silencing efficiency correlates with dsRNA length, dsRNA constructs of 400 nt to 500 nt and 800 nt were generated targeting single *FgCYP51* genes (CYPA-500/800, CYPB-400/800, CYPC-400/800). Additionally, the full-length cDNA of each *FgCYP51* gene (CYPA-full, CYPB-full, CYPC-full) was cloned without the start and the stop codon to avoid protein expression. The constructs were inserted into the vector p7U10-RNAi (Fig. S1) and transgenic *Arabidopsis* plants were generated. Resistance to *Fg* was analysed on detached leaves inoculated with 5×10^−4^ *Fg* conidia per ml and incubated at RT. At 5 days post inoculation (dpi) untransformed wt plants showed water-soaked spots with chlorotic and necrotic lesions representing typical symptoms of a successful *Fg* infection (Fig. 1A). In clear contrast, plants expressing CYP51-dsRNA of different length showed significantly reduced necrotic lesions compared to wt (Fig. 1A). There were no clear phenotypic differences between 400 nt and 800 nt or full-length constructs (Fig. 1A) and the reduction of the infection area was for nearly all constructs in a similar extent of about 50% to 60% in comparison to the control (Fig. 1B). The only exceptions were CYPB-800 that showed a higher resistance and reduced infection area by 77% whereas CYPA-800 showed the lowest resistance by reducing infection areas by only 34%. Concerning HIGS-mediated disease control, we found that there is no significant correlation between the length of the dsRNA precursor and the reduction of *Fg* infection on CYP51-dsRNA expressing *Arabidopsis* plants. Previously, we found that CYP51-dsRNA activity involved co-suppression in the respective non-targeted paralogous *FgCYP51* genes (Höfle et al., 2018 in revision). To analyse whether the observed phenotypes were provoked by co-silencing effects in the non-targeted *CYP51* genes, we measured the transcript levels of *FgCYP51* genes in the infected leaf tissue by qRT-PCR. As anticipated, the relative transcript levels of targeted genes *FgCYP51A, FgCYP51B*, and *FgCYP51C* were reduced after inoculation of leaves expressing the respective CYP-dsRNA constructs (Fig. 2). By comparing the 800 nt dsRNAs with the shorter 400 nt dsRNAs, the longer precursors showed a higher gene silencing efficiency and reduced the expression of all *FgCYP51* genes by 80% or more. For full-length constructs, only non-target gene expression could be determined because no gene specific primer that would not also bind in the original construct sequence was available for qRT-PCR. Silencing efficiency of non-target genes was high and over 60% in most cases. Notably, the strongest gene silencing efficiency was observed for dsRNAs of 800 nt in length (Fig. 2). Thus, CYPB-800 exhibited the strongest decrease in *Fg* infection of nearly 80% (Fig. 1).

**Fig. 1.**
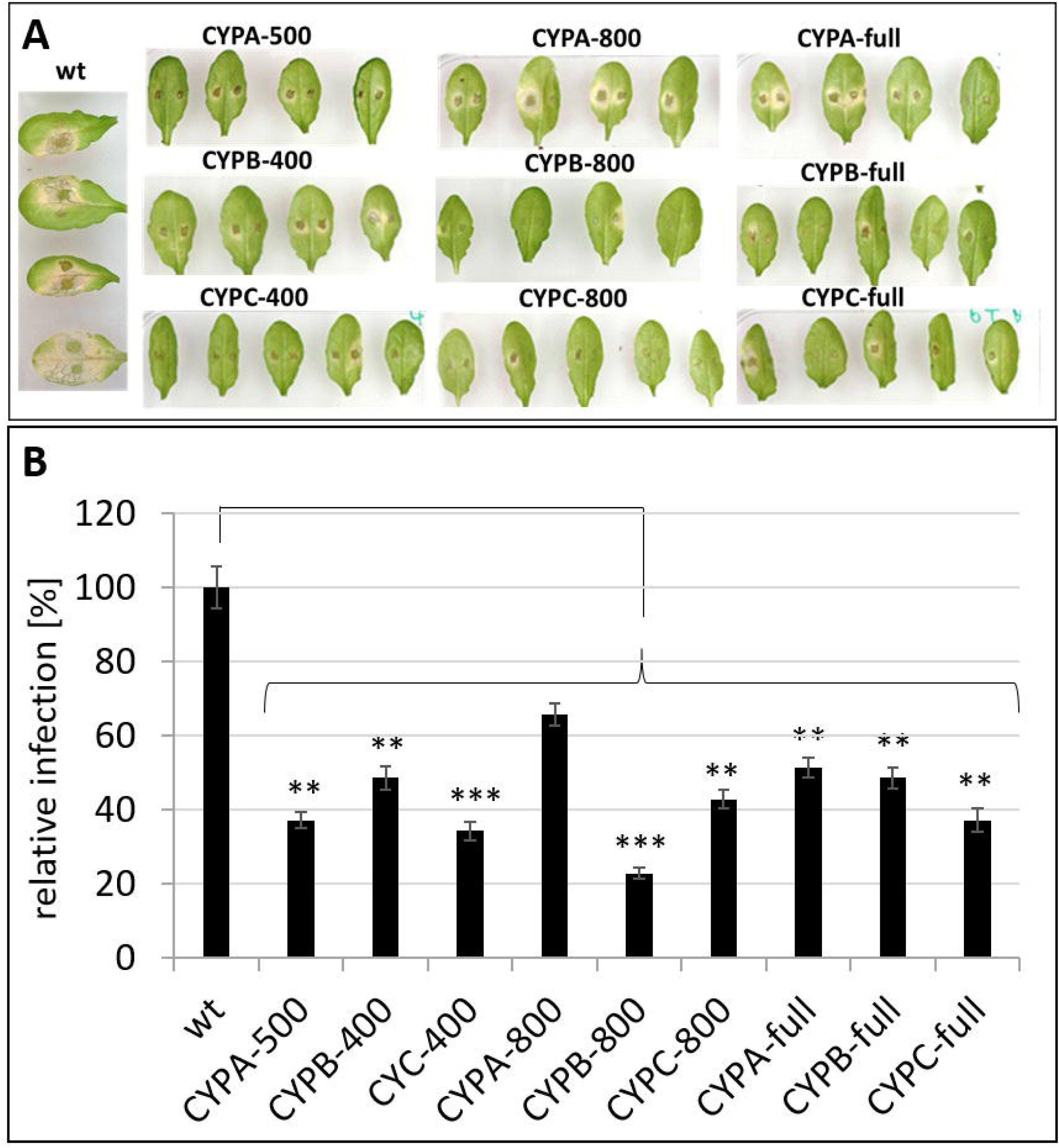
Host-Induced Gene Silencing in *Fg* on leaves of transgenic *Arabidopsis* expressing CYP51-dsRNAs of different lengths. **A**, fifteen detached rosette leaves of CYP51-dsRNA-expressing *Arabidopsis* plants (T2 generation) were drop-inoculated with 5 × 10^4^ conidia ml^-1^. Infection symptoms were evaluated at 5 dpi. **B**, quantification of the visibly infected area at 5 dpi shown as percent of the total leaf area. Error bars represent SE of two independent experiments each using 15 leaves of 10 different plants for each transgenic line. Asterisks indicate statistical significance (*p<0.05; **p<0.01; ***p < 0.001; students t-test).

**Fig. 2.**
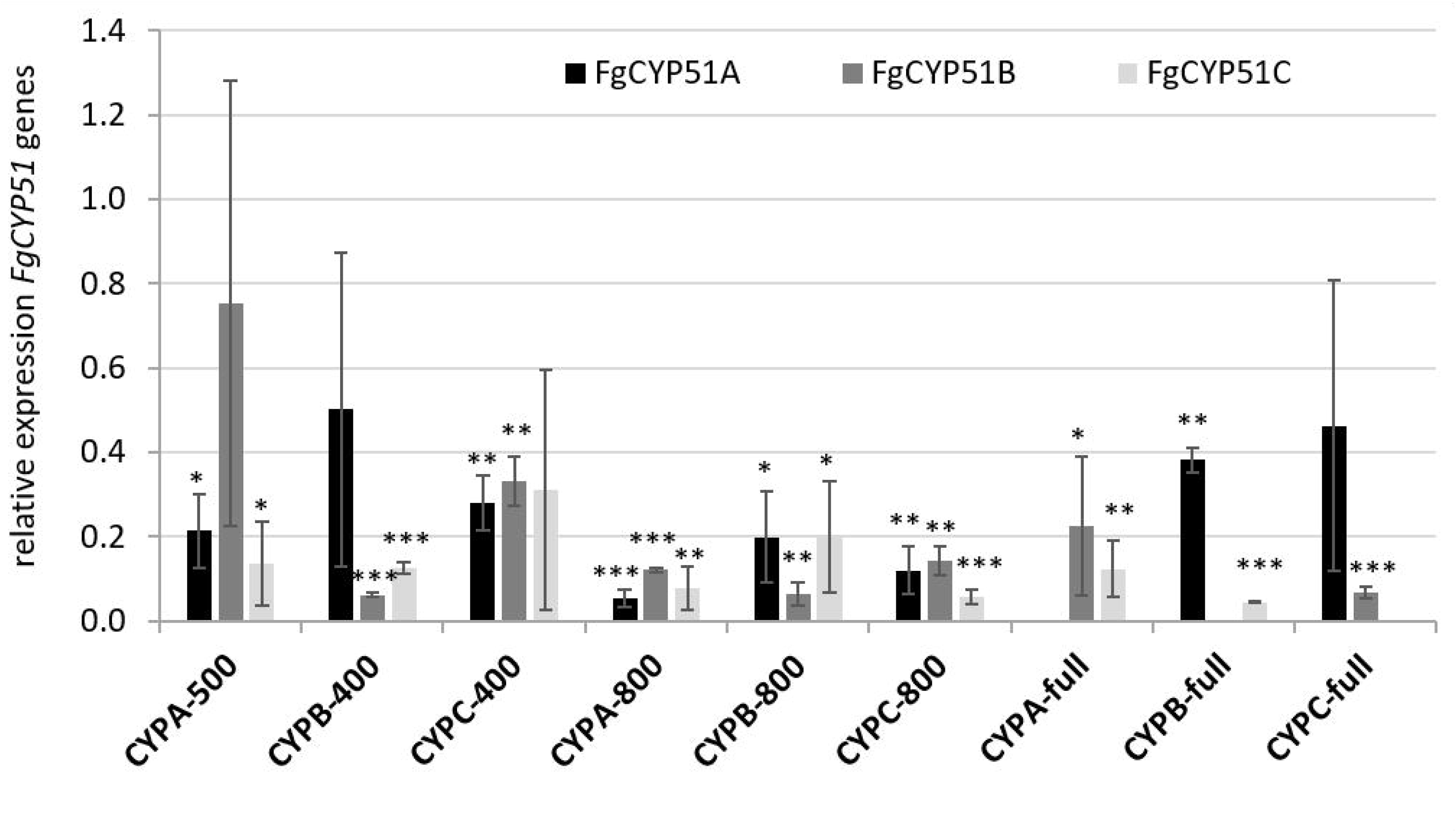
Silencing of *FgCYP51* genes of infected transgenic Arabidopsis leaves. Gene-specific expression of *FgCYP51A, FgCYP51B* and *FgCYP51C* was measured by qRT-PCR and normalized to fungal *EF1-α* (FGSG_08811) as reference gene. cDNA was generated after total RNA extraction from infected leaves at 5 dpi. The reduction in *CYP51* gene expression in the *Fg*-inoculated dsRNA-expressing leaves compared to the wt control was statistically significant. Error bars represent SD of two independent experiments each using 15 leaves of 10 different plants for each transgenic line. Asterisks indicate statistical significance (*p<0.05; **p<0.01; ***p < 0.001; students t-test).

Interestingly, CYPA-800 which had a similar silencing efficiency than CYPB-800, showed the lowest resistance by reducing infection areas by only 34% (Fig. 1). Consistent with this, we have recently shown that even strong silencing of *FcCYP51A* by almost 90% did not led to growth inhibition and morphological changes of *Fusarium culmorum* (*Fc*) *in vitro* cultures compared to only 40% silencing of *FcCYP51B* which resulted in 30% retardation of fungal growth as well as abnormal hyphal phenotypes (Koch et al. 2018). These results further support previous reports that have shown that *FgCYP51B* is the most important for ergosterol biosynthesis and thus survival of the fungus (Fan et al., 2013; Höfle et al., 2018 in revision; Liu et al., 2011; Machado et al., 2017; Koch et al., 2018)

By comparing the 800 nt dsRNAs with the shorter 400 nt dsRNAs, the longer precursors showed a higher gene silencing efficiency and reduced the expression of all *FgCYP51* genes by 80% or more (Fig. 2). Thus, further supporting our hypothesis that longer dsRNA led to a higher number of efficient siRNAs.

### Spray-induced gene silencing efficiency correlates with the length of the sprayed dsRNA

Spraying barley with dsRNA constructs of 200-300 nt in length targeting single *CYP51* genes of *Fg* was superior to HIGS-mediated *Fg* disease control (Höfle et al., 2018 in revision; Tab. 2). Encouraged by these findings, we assessed whether 400 nt, 800 nt as well as full-length CYP51-dsRNAs are also active in spray experiments. Detached barley leaves were sprayed with 20 ng μl^-1^ dsRNA and drop-inoculated 48 h later with a suspension of *Fg* conidia. After 5 dpi, necrotic lesions were visible at the inoculation sites of leaves sprayed with TE buffer (control). All CYP51-dsRNAs reduced the infection symptoms as revealed by significantly smaller lesions (Fig. 3A). We found strongest resistance after spraying with 400-500 nt constructs as the infection symptoms were reduced by 93% for CYPA-500, 89% for CYPB-400 and 90% for CYPC-400, respectively (Fig. 3B). Interestingly, infected areas of 800 nt constructs were reduced on average by 70% compared to the control (Fig. 3B), suggesting that the efficiency decreased by spraying 800 nt dsRNA. Recently, we have demonstrated that spraying a 791 nt long noncoding dsRNA (CYP3RNA), which targets the three *CYP51* genes *FgCYP51A, FgCYP51B* and *FgCYP51C* of *Fg* strongly inhibited fungal growth on barley leaves (Koch et al., 2016). Moreover, we have shown that SIGS was mediated by the uptake of the 791 nt dsRNA in the fungus and required the fungal RNAi machinery for the procession of siRNAs (Koch et al., 2016). Consistent with these findings, we found that 800 nt CYP51-dsRNAs inhibited *Fg* infection by 70% (Tab. 2). However, 800 nt dsRNAs targeting individual *FgCYP51* genes were less efficient compared to the 791 nt CYP3RNA that targets all three *FgCYP51* genes in parallel which led to 93% disease resistance (Höfle et al., 2018 in revision). This difference can be partly explained by co-silencing of *FgCYP51* genes.

**Fig. 3.**
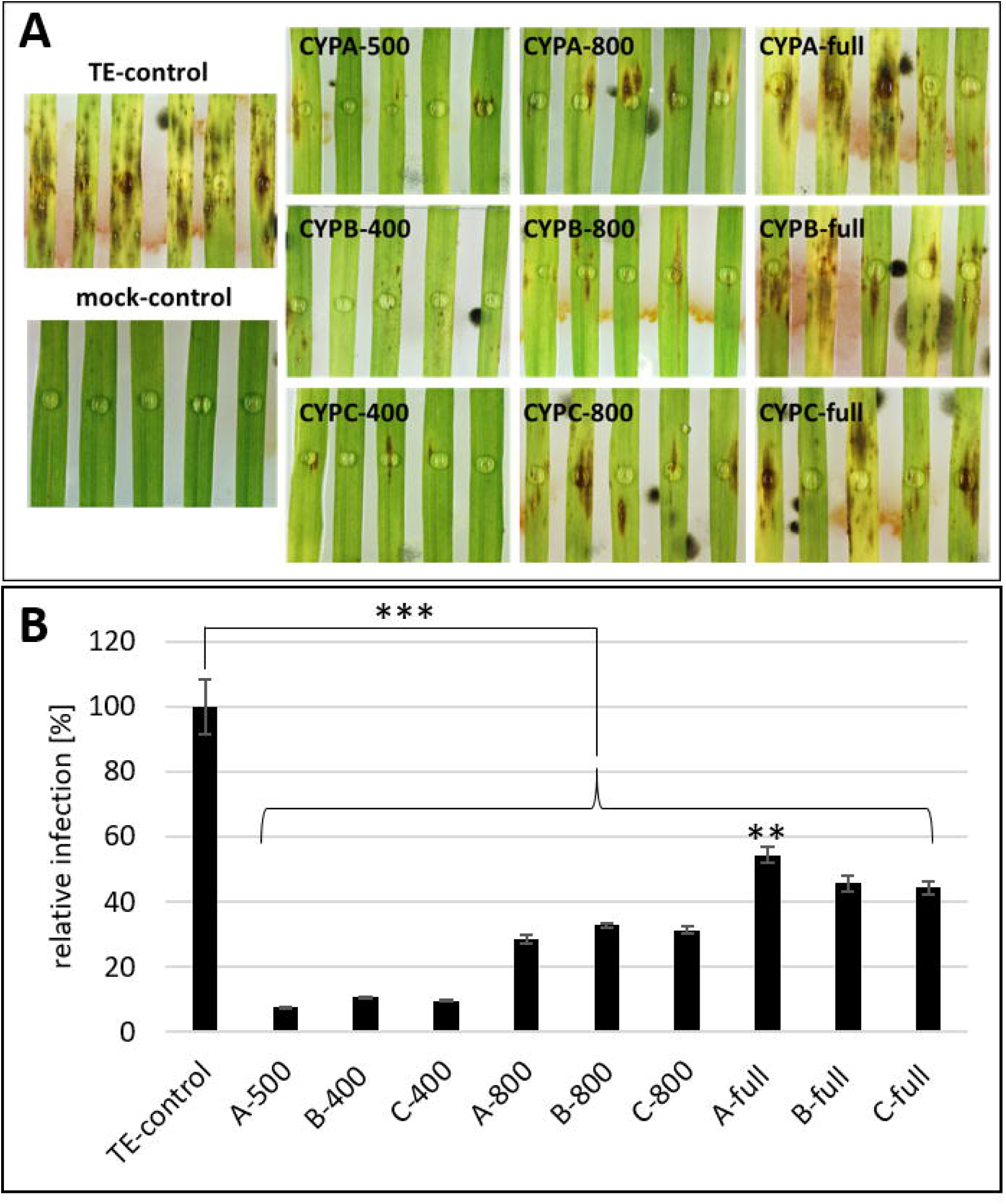
*Fg* infection of barley leaves that were spray treated with CYP51-dsRNAs of different lengths. **A**, Detached leaves of 3-week old barley plants were sprayed with CYP51-dsRNAs or TE buffer. After 48 h, leaves were drop-inoculated with 5 × 10^4^ conidia ml^-1^ and evaluated for infection symptoms at 5dpi. **B**, Infection area, shown as percent of the total leaf area for 10 leaves for each dsRNA and the TE control. Error bars indicate SE of two independent experiments. Asterisks indicate statistical significance (**p<0,01; ***p< 0,001; students t-test).

Previously we found that HIGS-as well as SIGS-mediated targeting of individual *FgCYP51* genes resulted in co-silencing of the non-targeted *FgCYP51* genes (Höfle et al., 2018 in revision). To further explore co-silencing effects, we calculated possible off-targets in *CYP51* genes for all tested CYP51-dsRNA constructs. Sequences of the different dsRNA constructs were split into k-mers of 18 bases and mapped to the coding sequences of the three *FgCYP51* genes, allowing a maximum of three possible mismatches. Based on these parameters, we calculated off-targets for all constructs in the respective non-target *FgCYP51* genes (Fig. 4) which is consistent with the observed gene silencing of all three *FgCYP51* genes (Fig. 2; Fig. 5).

**Fig. 4.**
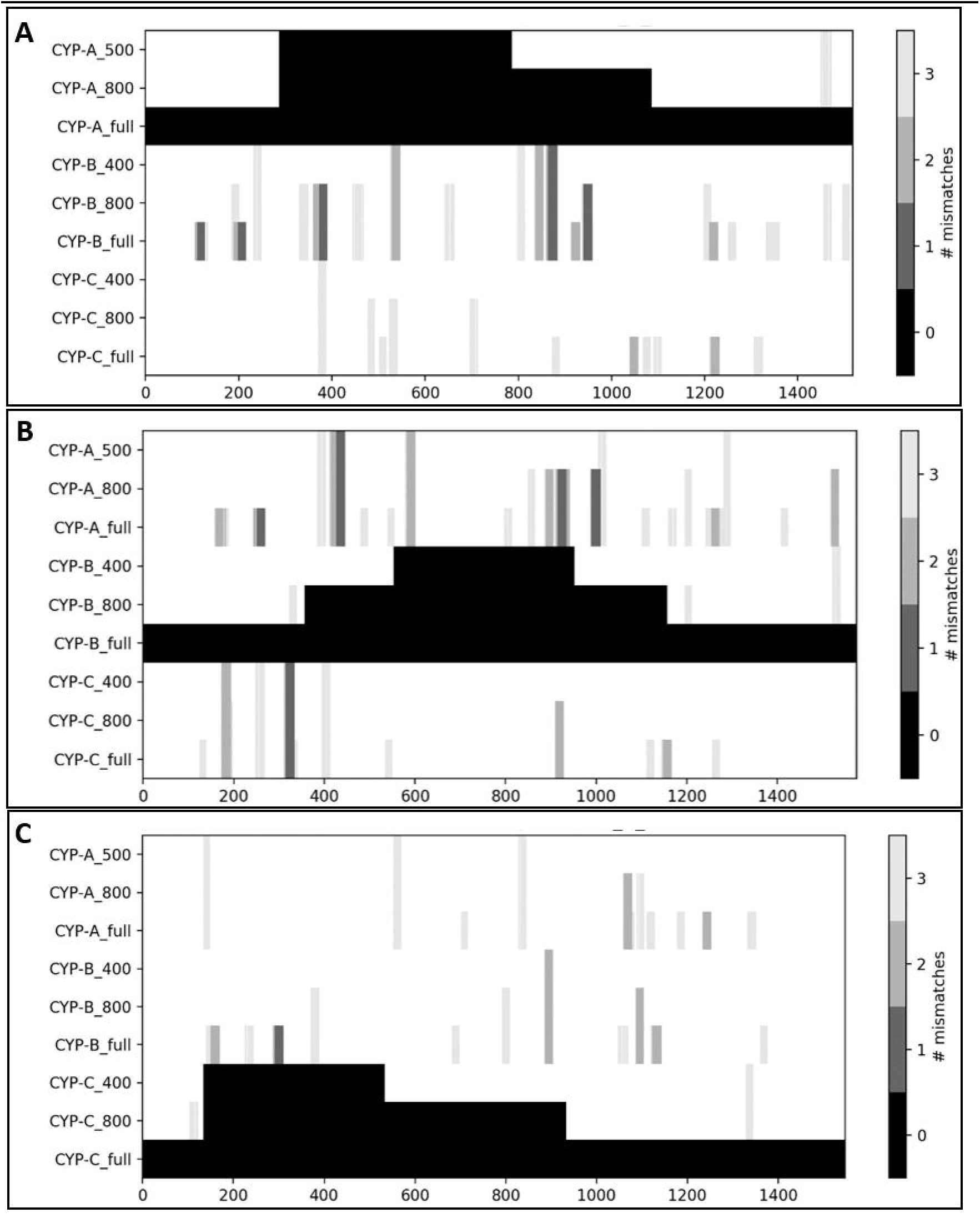
Off-target prediction for single CYP-dsRNA constructs with different length. Sequences of CYP51-dsRNAs (y-axis) were split into k-mers of 18 nt. These were mapped against the corresponding coding sequences (CDS) of *FgCYP51A* (**A**), *FgCYP51B* (**B**) and *FgCYP51C* (**C**). For each position within the CDS (x-axis) the k-mers that match with a specified number of mismatches is plotted.

**Fig. 5.**
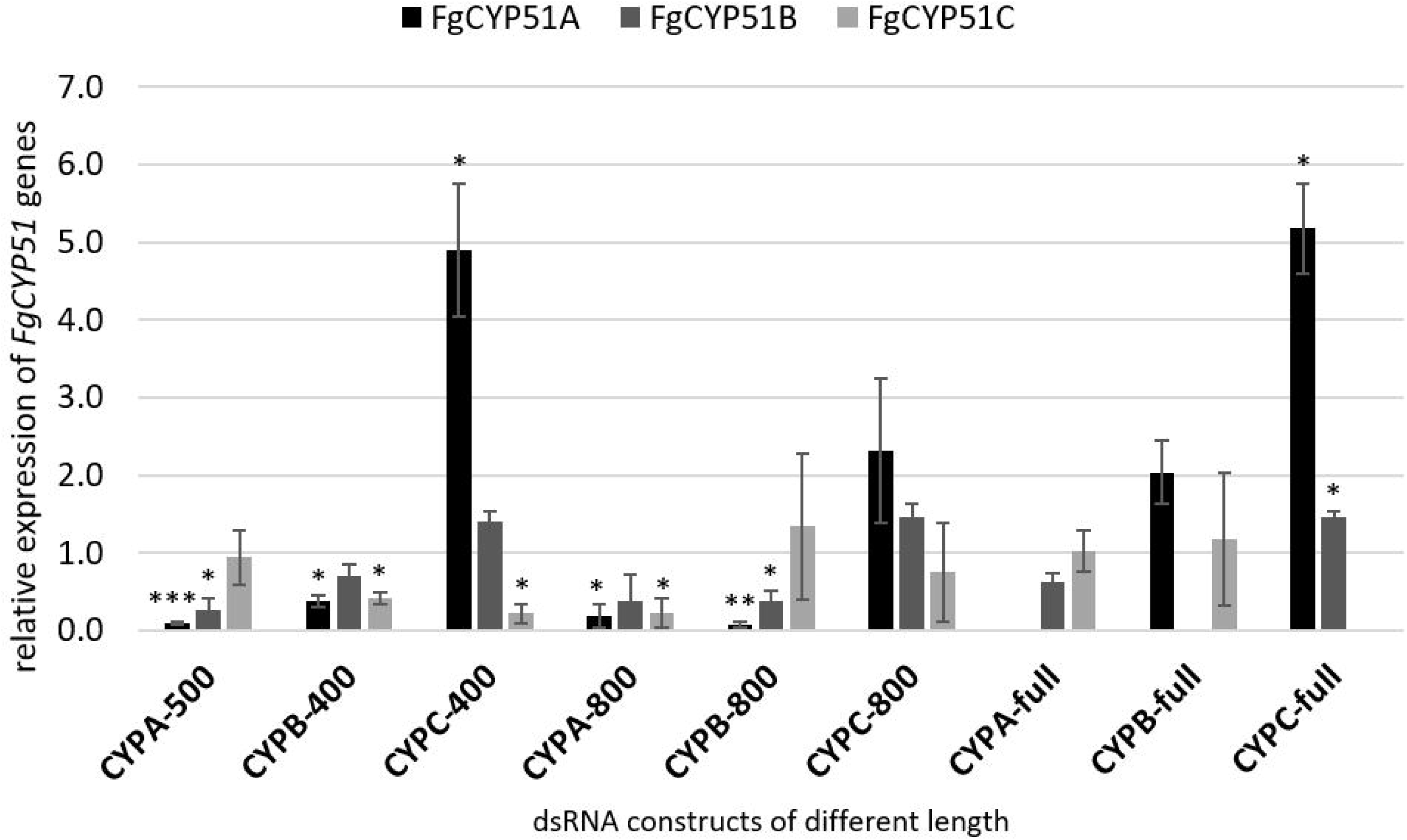
Silencing of *FgCYP51* genes of infected spray treated barley leaves. Gene-specific expression of *FgCYP51A, FgCYP51B* and *FgCYP51C* was measured by qRT-PCR and normalized to fungal EF1-α as reference gene. Detached leaves of 3-week-old barley plants were sprayed with CYP51-dsRNA or TE buffer. After 48 h leaves were drop inoculated with 5 × 10^4^ macroconidia ml^-1^. cDNA was generated at 5 dpi after total RNA extraction from infected leaves. Error bars represent SD of two independent experiments. Asterisks indicate statistical significance. (*p< 0,05; **p<0,01; ***p< 0,001; students t-test).

Consistent with our previous results, we observed that the number of off-targets increased with increasing length of the different CYP51-dsRNAs (Fig. 4). Thereby, we found that dsRNA constructs targeting *FgCYP51A* had the highest number of off-target hits in the non-targeted *FgCYP51B* and *FgCYP51C* genes (Fig. 4). This correlation resulted in the strongest silencing efficiencies for CYPA-500 and CYPA-800 dsRNA constructs (Fig. 5) thus, leading to the highest disease resistance efficiencies of 93% for CYPA-500 and 71% for CYPA-800 in SIGS, respectively (Tab. 2). Notably, *FgCYP51C*-derived constructs seem to have fewer off-targets in respective non-target genes than *FgCYP51A-* and *FgCYP51B*-derived constructs, reflected by qRT-PCR results (Fig. 5). Thereby the number of off-targets per construct increased with the length of the precursor RNA showing a maximum in the full-length constructs, as expected (Fig. 4). This was regardless of whether *FgCYP51A, FgCYP51B* or *FgCYP51C* was the actual target. Generally, *FgCYP51C*-derived constructs seem to have fewer off-targets in respective non-target genes than *FgCYP51A-* and *FgCYP51B*-derived constructs, although this was not reflected by qRT-PCR results (Fig. 2). To further prove whether longer dsRNAs result in higher numbers of siRNAs we predicted the number of siRNA hits for each CYP51-dsRNA constructs using SiFi (https://sourceforge.net/projects/sifi21) as prediction tool. Similar to what we observed for the off-target prediction we found a strong correlation between the length of the dsRNA precursor and the precursor-derived siRNAs (Tab. 1). However, this off-target based co-silencing effects were more obvious for SIGS than for HIGS (compare Fig. 2 with Fig. 5) further supporting our previous finding that SIGS involves the uptake of dsRNA by the fungus and requires the fungal RNAi machinery (Koch et al., 2016).

**Tab 1.**
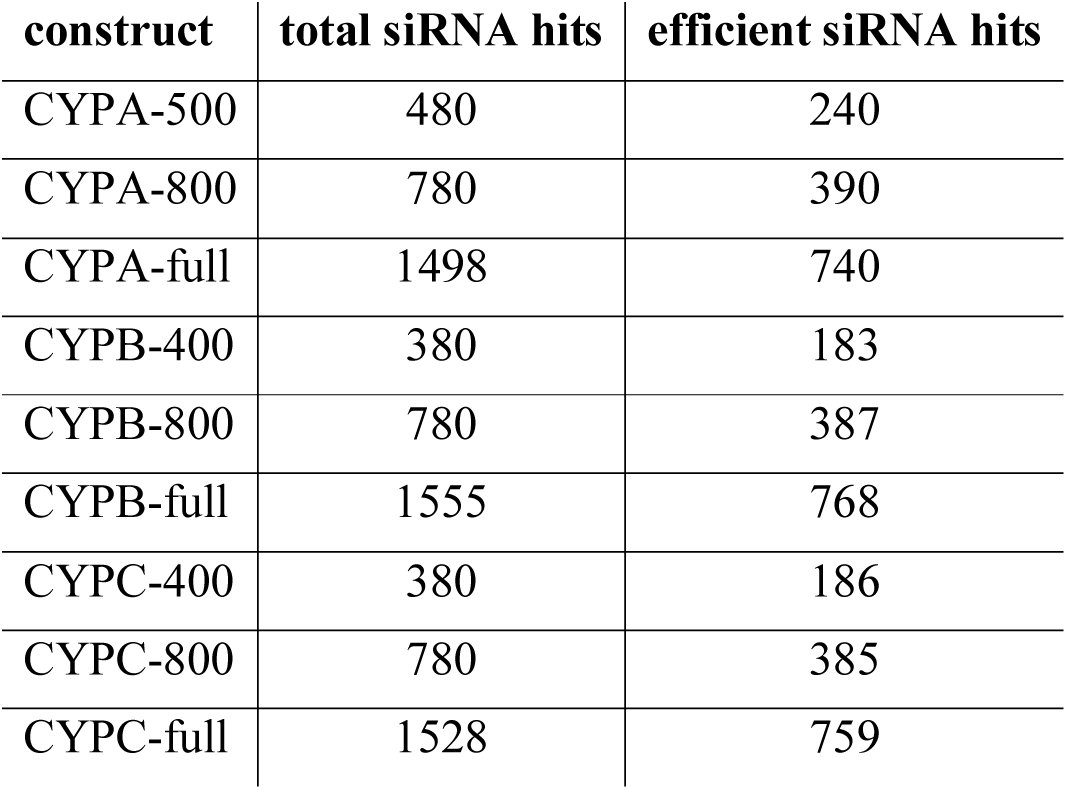
siRNA prediction of CYP51-dsRNA constructs using siFi21. Total and efficient siRNAs of 21 nt were predicted with the siFi21 software for the cDNA of the respective *FgCYP51* target genes *FgCYP51A, FgCYP51B* and *FgCYP51C*.

### Spray-induced gene silencing efficiency and uptake of dsRNA is limited by the length of the CYP51-dsRNA

Spraying with full-length CYP51-dsRNA resulted in the lowest decrease in *Fg* infection by at least 50% (Fig. 3B). Thus, we overserved a decrease in *Fg* infection that correlates with the length of the sprayed dsRNA (Tab. 2). However, expression analysis of *FgCYP51* genes in infected leaves showed target gene silencing and an overall strong co-silencing (Fig. 5). All constructs led to downregulation of respective non-targeted *CYP51* genes except for constructs targeting *FgCYP51C* (CYPC-400, CYPC-800 and CYPC-full), where an upregulation of the non-targeted *FgCYP51A* and *FgCYP51B* genes was measured (Fig. 5). This was consistent with the off-target prediction, where CYPC constructs showed the lowest number of siRNA that match to the *FgCYP51A* and *FgCYP51B* genes. Overall, these data show a strong correlation between resistance phenotypes induced by CYP51-dsRNA constructs and reduced expression of *CYP51* genes. Notably, we overserved that co-suppression was completely lost when leaves were sprayed with full-length CYP51-dsRNA (Fig. 5). Unfortunately, analysis of target gene silencing was not possible as there were no primers available that would not amplify the sprayed RNA as well.

**Tab 2.**
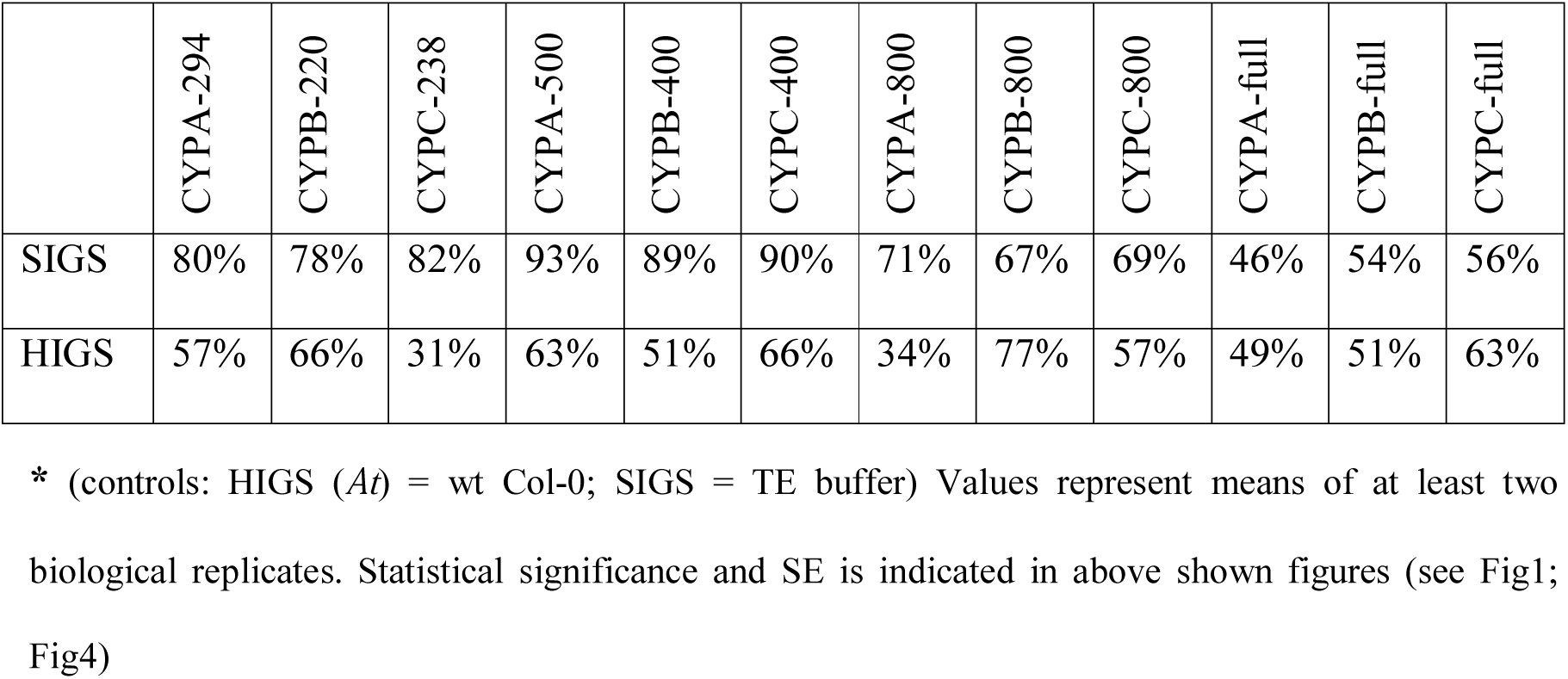
Growth inhibition of *Fg* during different RNAi-based silencing setups. Growth inhibition is shown as reduction in % of the infected leaf area in comparison to the control*.

However, the decrease in efficiency from 70% for 800 nt CYP51-dsRNAs to only 50% with the full-length dsRNAs indicate that the size of the dsRNA interferes with a sufficient uptake of dsRNAs by the fungus. To further prove this idea, we treated *Fg* with full length CYP51-dsRNAs *in vitro*. We found that there was no silencing of the non-targeted *FgCYP51* genes, indicating that *Fg* was not able to take up >1500 nt dsRNA from liquid cultures (Fig. S2). Unfortunately, we were not able to analyse target gene silencing directly as there were no primers available that would distinguish between dsRNA that was applied to the medium and silencing of *FgCYP51* target genes. However, as we observed 50% less infection symptoms after spraying barley leaves with full-length CYP51-dsRNA (Tab. 2) we anticipate that there are differences in the mechanism of fungal dsRNA uptake in SIGS compared to *in vitro* culture treatments. In *Aspergillus nidulans* as well as in *Aspergillus fumigatus* uptake of siRNAs resulted in gene silencing (Jöchl et al., 2014; Kalleda et al., 2013; Khatri and Rajam, 2007). In *Candida albicans* a structure dependent uptake efficiency could be observed as linear nucleic acids were taken up more efficiently than hairpin structures (Disney et al., 2003).

Congruent with the idea of different uptake/translocation mechanism we demonstrated that HIGS, even though 200-500 nt CYP51-dsRNA constructs were less efficient compared with SIGS, is not limited by the size of the dsRNA precursor the plants were transformed with. As dsRNAs are expressed *in planta* and subsequently processed by DCLs enzymes, we would expect that increasing the length of the dsRNA result in more siRNAs that were taken up by the fungus (Tab. 1). Unfortunately, and in contrast with our expectation, the data showed not such a correlation (Tab. 2). However, even for HIGS it must be considered that the host produced dsRNA precursor could be taken up and processed by the fungus itself. Consistent with our findings, a size dependent efficacy of dsRNA-mediated silencing of target genes has been observed in insects (Bolognesi et al., 2012; Saleh et al., 2006). Thereby the most efficient dsRNA length varies hardly among different species whereas most studies show successful gene silencing with dsRNA lengths from 140 to 520 nt (Andrade and Hunter, 2016). In the western corn rootworm silencing efficiency increased with increasing dsRNA lengths (60-200 nt) whereas a minimum of 60 nt dsRNA is required for successful gene silencing (Bolognesi et al., 2012). By targeting the same gene in aphid feeding studies, only dsRNA and not siRNAs were able to achieve gene silencing (Mulot et al., 2016). In *Leptinotarsa decemlineata* even a 1842 nt dsRNA resulted in mortality in feeding experiments (Baum et al., 2007; Huvenne and Smagghe, 2010). Interestingly, in the same study also shorter dsRNAs of 134 nt and 300 nt conferred gene silencing, suggesting that target gene selection is superior to dsRNA length.

## Conclusion

Previously, we demonstrated that gene silencing efficiency of individual *FgCYP51* genes depends on the number of functional siRNAs that reach the fungus (Koch et al. 2016). Given these results, questions on the design of new and potentially promising dsRNAs arise. Whereas bioinformatics provides a variety of tools for the design, the analysis, and the evaluation of sRNA agents, little is known about the influence of dsRNA design on target gene silencing efficiency. Therefore, we attempt to identify design principles of an optimal dsRNA trigger in order to improve/guarantee efficacy and specificity of HIGS- and SIGS-based plant protection approaches. By comparing efficiencies of infection reduction of HIGS with SIGS we observed a strong bias between those two approaches (Tab. 2). For example, CYP51-dsRNA construct CYPC-238 (Höfle et al., 2018 in revision; Tab. 2) exhibited 78% reduction of infection in SIGS compared to only 31% for HIGS (Tab. 2). Another example is CYPA-500 which showed 93% efficiency in SIGS but only 63% decrease in *Fg* infection in HIGS. In other words, for dsRNA constructs of 200-500 nt HIGS is around 30% less efficient compared with SIGS (under lab conditions) (Tab. 2). However, we observed that with increasing length of the CYP51-dsRNA construct differences in efficiencies between HIGS and SIGS approaches became less obvious. If we compared CYPB-800 and CYPC-800 dsRNA constructs, SIGS was only 10% more efficient than HIGS (Tab. 2). Notably, if we than compared efficiencies of CYP51-dsRNA constructs that were generated out of the full-length sequence (>1500 nt) of the individual *FgCYP51* genes, HIGS and SIGS showed the same level of around 50% reduction of *Fg* infection (Tab. 2). Importantly and in clear contrast to HIGS, we measured a decrease in SIGS-mediated *Fg* disease resistance that is probably correlated with the length of the dsRNA construct that was sprayed. More explicated, SIGS-based efficiencies decreased from 90% for 200-500 nt constructs to 70% for 800 nt constructs to 50% for >1500 nt dsRNA constructs.

Taken together our results strongly support the notion that uptake of dsRNA of 200-800 nt in length and procession of siRNA by the fungus is more efficient in terms of disease control than uptake of siRNA via HIGS. However, little is known about how and which silencing signals (i.e. siRNAs and/or dsRNA precursors) are transferred from plant into fungal cells. Thus, since we have no evident proof for our conclusion this statement remains speculative. Therefore, further research must address the question: What is the optimal dsRNA design for uptake, translocation and silencing efficiency of the RNAi trigger compound?

## Methods

### Construction of CYP51 containing p7U10 RNAi vectors

CYP51-dsRNA constructs CYPA-500/800/full, CYPB-400/800/full and CYPC-400/800/full were amplified from *Fusarium graminearum* IFA65 cDNA using gene specific primer (Tab. S1) and inserted into the HindIII and XmaI restriction sites of p7U10 RNAi (Fig. S1).

### Generation of transgenic *Arabidopsis thaliana* plants

p7U10 plasmids for transformation of *Arabidopsis* were introduced into the *A. tumefaciens* strain AGL1 by electroporation. Transformation of *Arabidopsis* was performed with the floral dip method as described (Bechtold and Pelletier, 1998) and transgenic plants were selected on ½ MS agar plates containing BASTA (7 μg/ml).

### Plant infection assays and spray application of dsRNA

Fg IFA65 was grown on SNA agar plates at 22°C in an incubator (BINDER). For all leaf inoculation assays, Fg-IFA65 conidia concentration was adjusted to 5 × 10^4^ macroconidia ml^-1^ in ddH20 containing 0.002% Tween-20. After inoculation, plates were stored at RT and infection symptoms were assessed at 5 dpi. To evaluate infection severity, fungal growth was determined by measuring the size of chlorotic and necrotic lesions using the ImageJ software (https://imagej.nih.gov/ij/index.html).

For the *Arabidopsis* – *Fusarium* infection, 15 rosette leaves of ten different 5-wk-old plants of each transgenic line and control plants [Col-0 wild-type (wt)] were detached and transferred in square petri dishes containing 1% agar. Inoculation of *Arabidopsis* was done by wound inoculation of detached leaves with 5 μl *Fusarium* conidia suspension on each leaf side. Wounding was performed by scratching of the leave surface with a pipette tip. At 5 dpi leaves were frozen in liquid nitrogen and subjected to RNA extraction and cDNA synthesis.

For spray application, dsRNA was generated using MEGAscript RNAi Kit (Invitrogen) following the manufacturer’s instructions. p7U10 plasmids containing CYP51-dsRNA constructs were used as template. Primer pairs with T7 promoter sequences at the 5’end of both forward and reverse primers were designed for amplification of dsRNA (Tab. S1). The dsRNA, eluted in TE-Buffer (10 mM Tris-HCl pH 8.0, 1 mM EDTA), was diluted in 500 μl water to a final concentration of 20 ng μl-1. For the TE-control, TE-buffer was diluted in 500 μl water corresponding to the amount that has been used for dilution of the dsRNA. Typical RNA concentration after elution was 500 ng μl-1, representing a buffer concentration of 400 μM Tris-HCL and 40 μM EDTA in the final dilution. Detached barley leaves were covered before spraying with a plastic tray leaving only the upper part (approximately 1 cm) uncovered. After spraying, dishes were kept open until the surface of each leaf was dried. After 48 h, leaves were drop-inoculated as described above.

### Quantification of fungal transcripts by Quantitative Real-Time PCR (qRT-PCR)

Before cDNA synthesis, remaining DNA was digested by DNAse I (Thermo Scientific) using RiboLock RNAse Inhibitor (Thermo Scientific) for 30 min at 37°C. For cDNA synthesis 1 μg digested RNA was used. cDNA synthesis was performed using qScriptTM cDNA synthesis kit (Quanta). Quantitative Real-Time PCR (qRT-PCR) was performed with freshly synthetized cDNA in the QuantStudio 5 Real-Time PCR system (Applied Biosystems) in 384-well plates using SYBR® green JumpStart Taq ReadyMix (Sigma-Aldrich). For each sample three replicates were performed, and target transcript levels were determined using gene specific primer (Tab. S1) via the 2^-Δ Δ Ct^ method (Livak and Schmittgen, 2001) by normalizing the amount of target transcript to the amount of reference transcript.

### Bioinformatic off-target analysis

The precursor sequences of CYP51-dsRNAs were split into k-mers of 18 bases. These sequences were mapped to the coding sequences of *FgCYP51* genes (CDS) of *Fusarium graminearum* strain PH-1 (GCA_000240135.3) with Segemehl (Hoffmann et al., 2009) using the following settings: accuracy of 60, report all targets, max seed distance of 4, max e-value of 20. The hits were filtered for an edit distance of 0, 1, 2 and 3. For each sequence the mapping depth per position and edit distance was plotted with Matplotlib (Hunter, 2007; Michael DroettboomNIH et al., 2017). The results were reported as plots. The analysis is implemented as an internal pipeline using Nextflow (Di Tommaso et al., 2017).

---

**Fig. S1.**
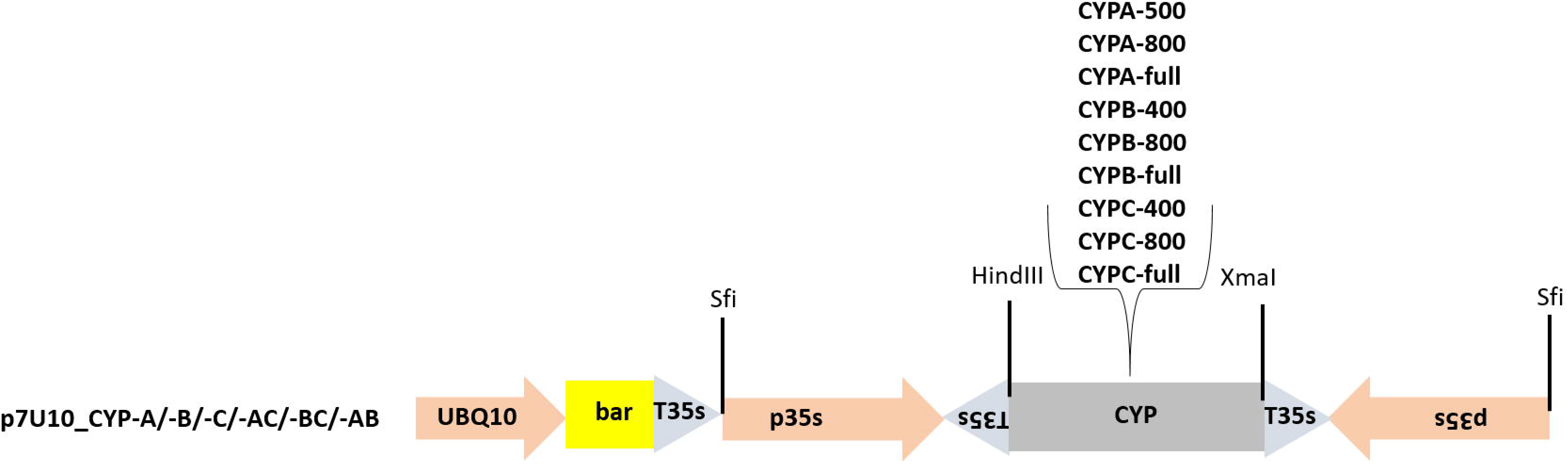
Schematic representation of RNAi vectors CYPA-500, CYPB-400, CYPC-400, CYPA-800, CYPB-800, CYPC-800, CYPA-full, CYPB-full and CYPC-full used for transformation of Arabidopsis.

**Fig. S2.**
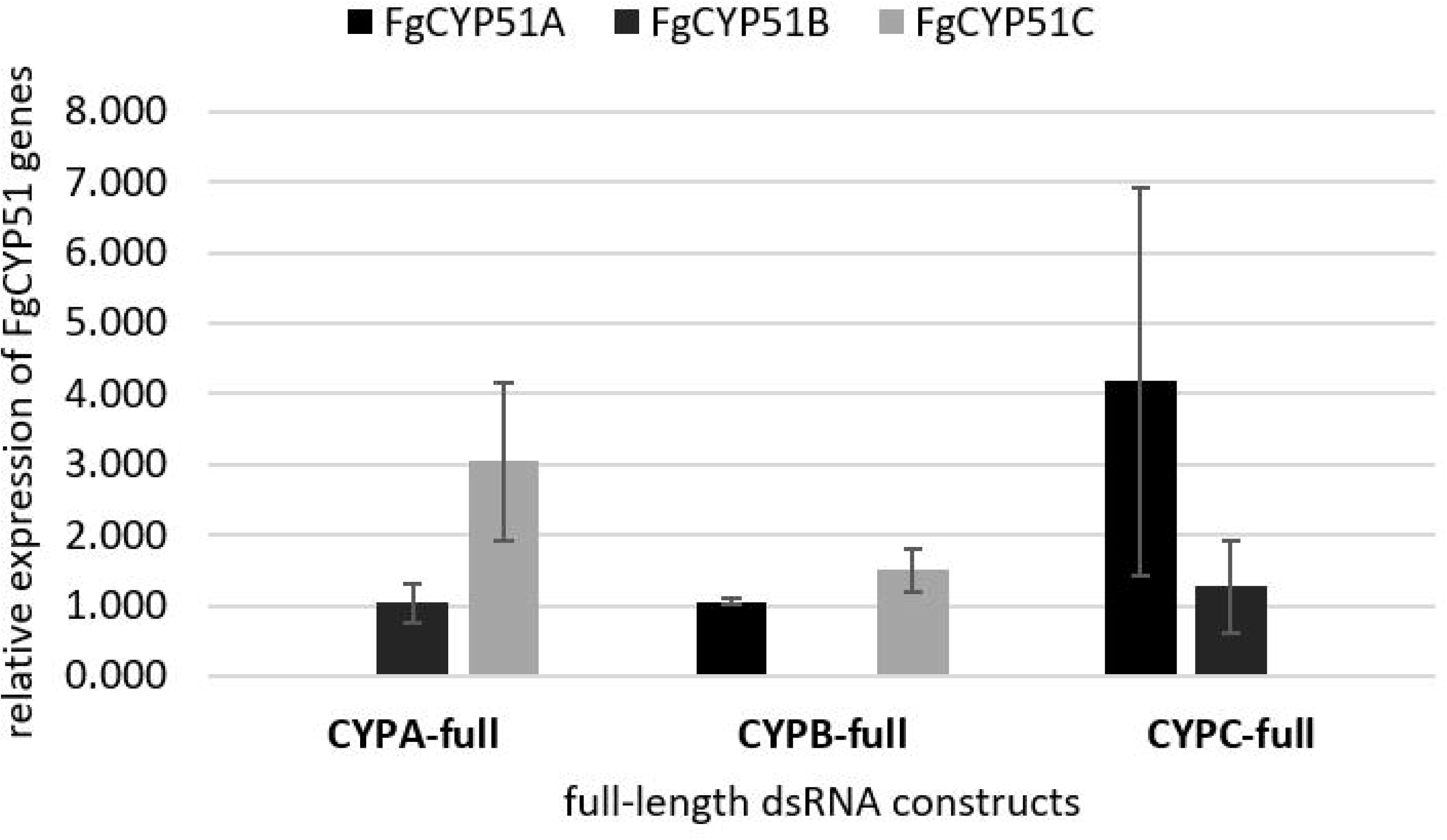
Silencing of *FgCYP51* genes in *Fg in vitro* cultures after treatment with full-length CYP51-dsRNA constructs. A conidia suspension (100 μl) containing 150,000 conidia ml^-1^ of *Fg* in SNA was incubated in 96-well plates for 2 days at RT with 2 μg dsRNA. cDNA was generated after total RNA extraction from *in vitro* cultures. Gene-specific expression of *FgCYP51A, FgCYP51B* and *FgCYP51C* was measured by qRT-PCR and normalized to fungal *EF1-α* (FGSG_08811) as reference gene. Error bars represent SE of three independent experiments. Asterisks indicate statistical significance (*p<0,05; **p<0,01; ***p < 0,001; students t-test).

**Tab S1.**
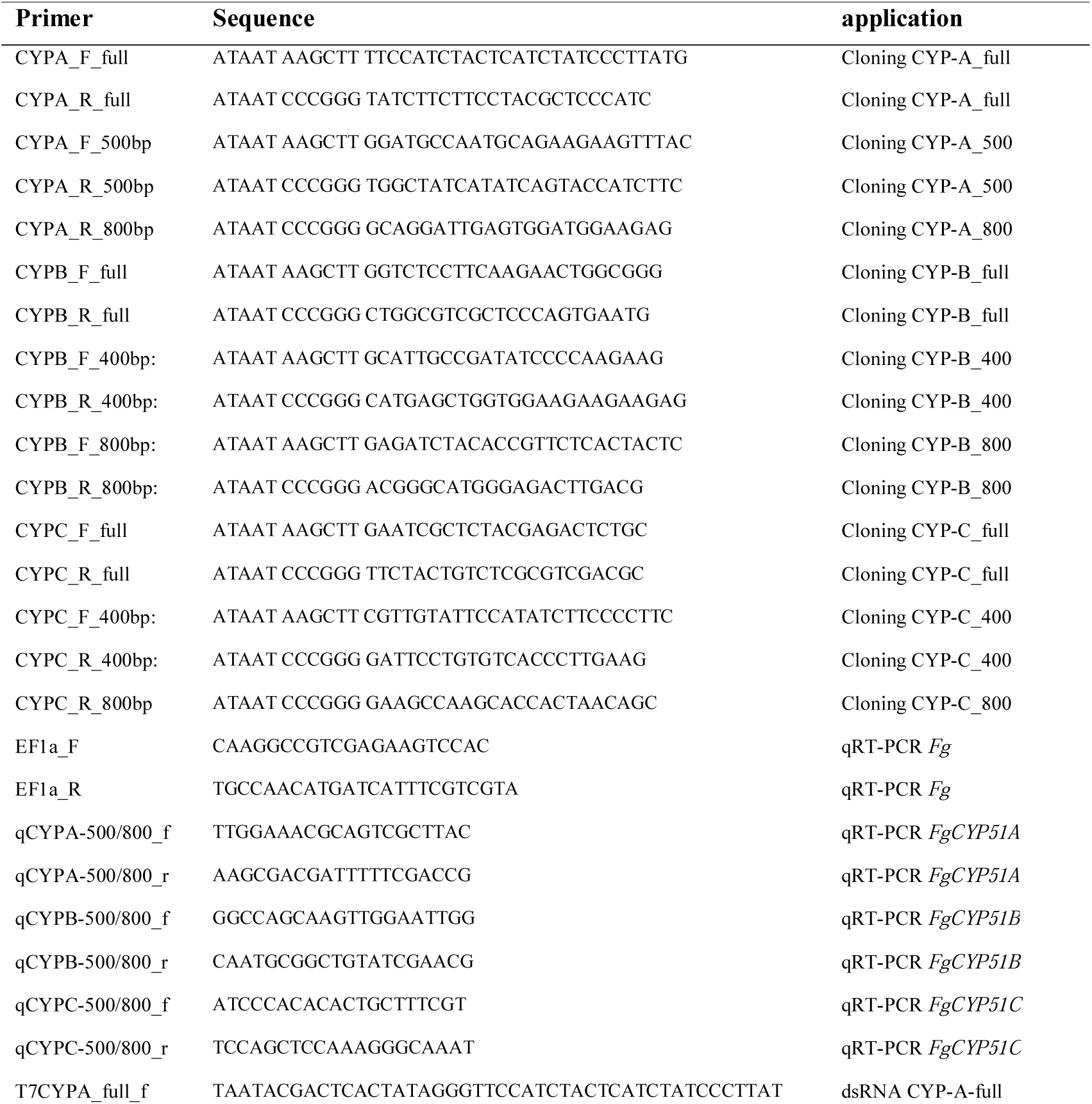
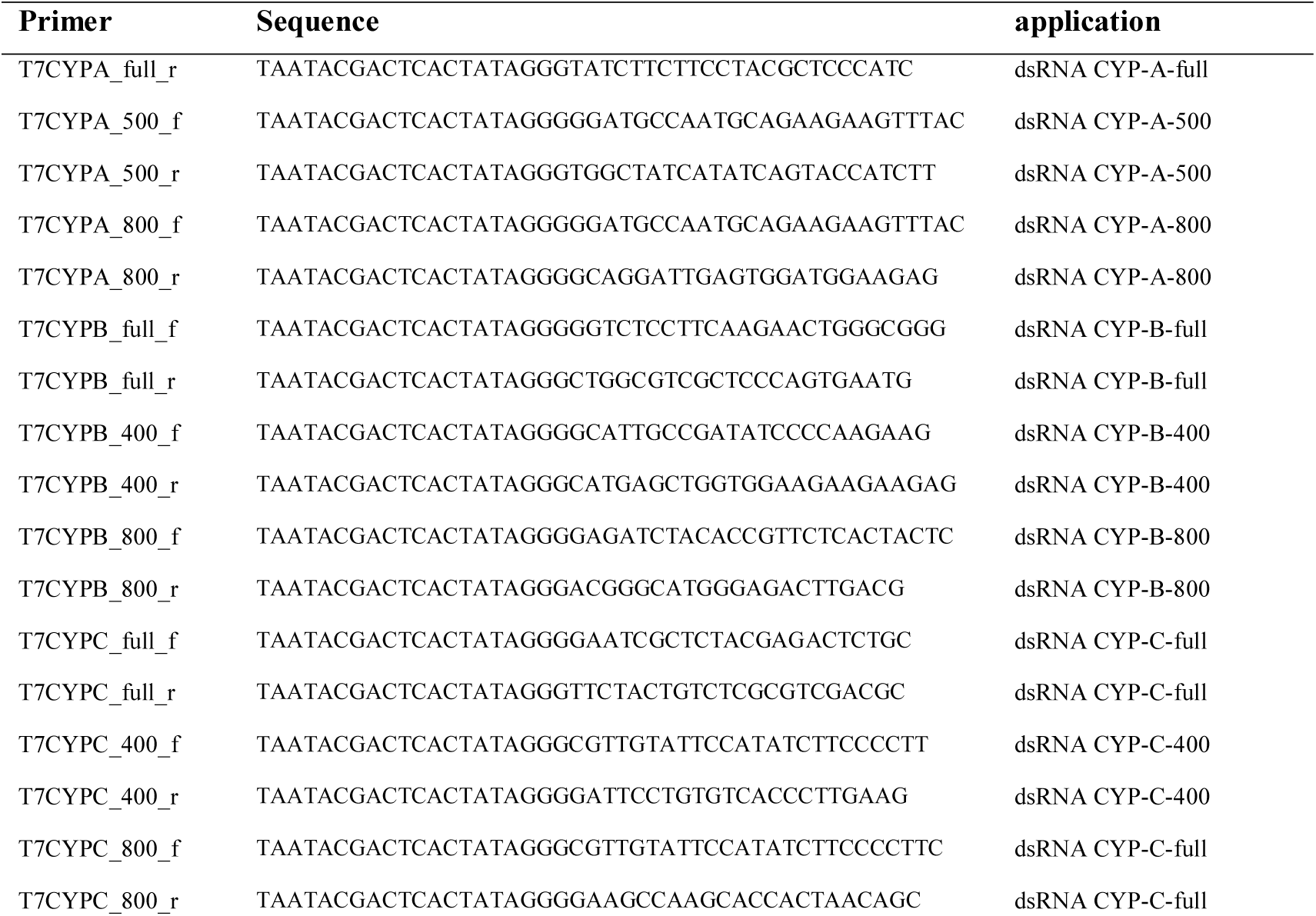
Primers used in this study for generation of CYP51 RNAi silencing constructs and qRT-PCR.

## Acknowledgements

We thank D. Biedenkopf for excellent technical assistance. C. Birkenstock, U. Schnepp and V. Weisel for excellent plant cultivation. This work was supported by Deutsche Forschungsgemeinschaft to AK.

## Competing financial interests

The authors declare no competing financial interests. Work on Fuarium CYP3RNA (Koch et al. 2013) is subject of an patent application (WO2015004174A1).

## Author Contributions

A.K. and L.H. wrote the manuscript; L.H. and A.K. designed the study; L.H. and A.S. conducted the experiments; A.K. and L.H. analyzed all data and drafted the figures. L.J. performed bioinformatics analysis. All authors reviewed the final manuscript.

